# A Comprehensive Meta-Analysis of Breast Cancer Gene Expression

**DOI:** 10.1101/2024.08.30.610515

**Authors:** Ifeanyichukwu O. Nwosu, Stephen R. Piccolo

## Abstract

**Background:** Triple-negative breast cancers (TNBC) occur more frequently in African Americans and are associated with worse outcomes when compared to other subtypes of breast cancer. These cancers lack expression of estrogen receptor (ER), progesterone receptor (PR) and human epidermal growth factor receptor 2 (HER2) and have limited treatment options. To shed light on mechanisms behind these differences and suggest novel treatments, we used a meta-analytic approach to identify gene expression differences in breast tumors for people with self-reported African or European ancestry; additionally, we compared gene expression levels based on ER, PR, HER2 and TNBC status.

**Methods:** After gathering and standardizing gene expression data and metadata from 106 datasets (representing 27,000 samples), we identified genes that were expressed differently between these groups via random-effects meta-analyses. To evaluate the robustness of these gene lists, we devised a novel computational methodology that uses cross validation and classification. We also computed overlaps between the most significant genes and known signaling pathways.

**Results:** Using a false discovery rate threshold of 0.05, we identified genes that are known to play a significant role in their respective breast cancer subtypes (e.g., *ESR1* for ER status and *ERBB2* for HER2 status), thus confirming the validity of our findings. We also discovered genes that have not been reported previously and may be new targets for breast cancer therapy. *GATA3*, *CA12*, *TBC1D9*, *XBP1* and *FOXA1* were among the most significant genes for ER, PR, and TNBC. However, none of these genes overlapped with HER2 status, supporting prior research that HER2 tumors are mechanistically different from endocrine breast cancers. The genes identified from the race meta-analysis—including *DNAJC15*, *HLA-DPA1*, *STAP2*, *CEP68*, *MOGS*—have not been associated previously with race-specific breast-cancer outcomes, highlighting a potential area of further research.

**Conclusions:** We have carried out a large meta-analysis of breast cancer gene expression data, identifying novel genes that may serve as potential biomarkers for breast cancer in diverse populations. We have also developed a computational method that identifies gene sets small enough to be analyzed and explored in future studies. This method has the potential to be applied to other cancers.

## Background

Cancers arise as a result of alterations in groups of genes known as proto-oncogenes and tumor suppressor genes^1,2^. These definitions cover hundreds of genes^3,4^ that have been implicated in tumorigenic processes, with an average of 33 to 66 mutations found in solid tumors^5^. This represents an enormous amount of genetic variation, even within the same tissue types^6^, making it difficult to develop a “one size fits all” approach to treatments, or to understanding cancer biology in general. Among cancers, breast cancer is the most common type of cancer in women worldwide, and accounts for 12.5% of new cancer cases in the world each year^7,8^. It is also the second most common cause of cancer deaths among women in the U.S. In 2023, the American Cancer Society estimated 297,790 new cases of invasive breast cancer diagnosed in women in the US alone^9^. This is an enormous public health burden. Advances in treatment, coupled with early diagnoses, have improved survival rates and lowered mortality^10,11^ for breast cancer patients. However, researchers continue to search for methods to assign the most appropriate types of therapies^12,13^ and thus to tailor treatments to patients.

Despite these advancements, the heterogeneous nature of the disease^14^ and the complex genetic landscape^15^ still make it difficult to unravel the underlying molecular mechanisms governing its pathogenesis and progression. Furthermore, there exist racial disparities among population groups^16^. For example, the American Cancer Society reports that survival rates for all cancers are lower for individuals of African descent when compared to those of Caucasian descent^17^. These differences are largely due to cancers of the breast in women, but this is also true for prostate^18^ and lung^19^ cancers. Various studies^20–22^ have highlighted the multifactorial causes of these disparities. These include socioeconomic factors such as access to healthcare, income levels, as well as treatment delays and trust in physicians. However, these factors do not fully explain these disparities^23,24^. The question thus arises as to factors potentially driven by biological causes.

Gene expression is the process by which the information encoded in genes is used to direct the synthesis of proteins via RNA transcripts. It is assessed by measuring the number of RNA transcripts for each gene in a tissue sample; the sum of a sample’s RNA transcripts is referred to as its transcriptome. High-throughput genomic technologies have enabled researchers to produce huge amounts of gene-expression data^25–27^. The analysis of gene expression data is a powerful tool for deciphering molecular signatures associated with different breast cancer subtypes^28^, predicting patient prognoses^29^ and guiding therapeutic decision-making^30^. Extensive research has shown that studying the transcriptome can help to identify functional mechanisms behind biological processes. This is validated by a number of studies which have explored gene expression patterns in cancer; these have found patterns associated with race^31–34^ and identified connections between race and hormone receptor status^35,36^. These studies, however, have used a limited number of samples (between 52 and 627 per study). Some of these datasets are publicly available, while others are not.

Traditionally, clinical subtypes of breast cancer have been defined based on a patient’s immunohistochemistry (IHC) profile. These profiles are determined by the combined expression status of three receptors: estrogen receptor (ER), progesterone receptor (PR), and human epidermal growth factor receptor 2 (HER2)^37^. It has also been observed that women of African descent more commonly have tumors negative for ER, PR and HER2^38^. These cancers are typically referred to as triple negative breast cancers (TNBC). TNBCs are associated with worse outcomes when compared to other subtypes^39^ and occur more frequently in African Americans^40,41^ than other groups. Some studies have identified biological processes that may contribute to enhanced disease aggressiveness in African-American patients, including angiogenesis, chemotaxis^42^, and the insulin-like growth factor 1 (IGF1) pathway^43^.

Building on a large curation effort that we performed with publicly available data^44^, we identified gene-expression datasets that include race metadata and/or metadata for ER, PR and HER2 status. In total, we have data for over 27,000 samples across 47 datasets with ER status, 35 datasets with PR status, 28 datasets with ER status and 7 datasets with race information. With these datasets, we applied a meta-analytic method that combines data from multiple independent scientific studies, with the aim of identifying consistent patterns^45^. Our research questions can be broadly divided into: A) which genes are most consistently expressed differently between individuals who self-identify as having European ancestry or African ancestry, B) which genes are consistently differentially expressed between individuals who are positive or negative for ER, PR, HER2, or TNBC status, and C) which signaling pathways are associated with these genes. By addressing these questions, we aim to uncover molecular mechanisms underlying breast cancer disparities and hormone receptor statuses, potentially informing personalized treatment strategies and improving outcomes across diverse populations.

## Methods

### Data Acquisition

In a prior study^44^, we gathered 114 gene expression datasets from public repositories including Gene Expression Omnibus^46,47^, ArrayExpress^48^, the Molecular Taxonomy of Breast Cancer International Consortium (METABRIC) project^49^, TCGA^50^ and the SCAN-B Initiative^51,52^ in a bid to enable researchers to address a variety of questions related to breast-cancer transcriptomics. For this study, we used a subset of those datasets (Additional File 1). We identified datasets for which one or more of the following metadata variables were available: self-reported race/ancestry, ER status, PR status, and/or HER2 status.

### Data Preparation

For the comparison based on self-reported ancestry, we filtered the data to focus on people described as “White”, “Caucasian” or “European American” and people described as “Black” or “African American”. We focused on these groups because they were most common in the data and because an understanding of disparities between these groups is critically needed. We removed samples that did not fall into either of these categories. Next, we standardized the metadata variable names because different studies used different terms to represent the same variables. Samples that were labeled “Caucasian”, “W”, “w”, “European American”, or “white” were all categorized as “White”, while samples labeled “African American”, “B”, “b”, “Black or African American”, or “black” were categorized as “Black”. For the cell-surface receptors (ER, PR and HER2), we standardized each value as “positive” or “negative,” as determined by the original authors. We excluded samples with missing values for these variables. For ambiguous data where we could not readily identify what the values referred to, we referred to the journal article(s) associated with each dataset. If we found more information to clarify what those values meant, we standardized and included them in our study; otherwise, we filtered them out. For example, in one study, we found that the values “0/1”, “0-1”, “1+”, and “1+ (only 1 core pos.)” referred to HER2 negative samples. When determining TNBC status, we implemented logic to handle missing values. We marked a tumor as TNBC-negative only if it was known that the tumor was negative for all three markers (no missing data). If a tumor was positive for any marker, we inferred that the tumor was *not* TNBC-negative, even if other markers were missing. Additionally, we identified tumors for which race and TNBC status were available and performed a comparison between individuals of African or European ancestry.

For each comparison (Black and White, ER+ vs ER-, PR+ vs PR-, HER2+ vs HER2-, TNBC vs non-TNBC, and Black TNBC vs White TNBC), we selected datasets that had at least 10 samples in each group. For example, when comparing ER status, we required a minimum of 10 “ER+” samples and 10 “ER-” samples. We also determined the relative proportion of the groups and ensured the ratio between groups was no larger than 20:1 (in either direction). We did this to prevent our results from being skewed due to class imbalance.

### Meta-analysis

Different gene-expression profiling platforms and annotations produce data for different genes. Therefore, for each comparison, we first selected genes that were common across all datasets being used for that comparison. There were 11,049 common genes for race, 10,809 common genes for ER status, 10,960 common genes for PR status, 10,926 common genes for HER2 status, 10,960 for triple negative status, and 11,049 for race + TNBC.

We carried out the meta-analyses using the DExMA^53^ package (version 1.10.7) in R^54^(version 4.3.2). We used the following parameters:

1. typeMethod: random effects model;
2. missAllow: The maximum proportion of missing values allowed in a sample = 10%;
3. proportionData: The minimum proportion of datasets in which a gene must be present = 90%.

After the meta-analysis was completed, we filtered the results based on False Discovery Rate (FDR) values less than 0.05 and arranged them in descending order based on the absolute value of the effect size. This ensured that the top-ranked genes were both statistically significant and had a relatively large effect size.

Meta-analyses commonly result in thousands of statistically significant genes. It is difficult to interpret such large lists. We wished to estimate the number of significant genes that would effectively discriminate between the groups while being as parsimonious as possible. Furthermore, Venet et al^55^ discuss that empirically identified gene-expression signatures are sometimes no more predictive of breast-cancer outcomes than random gene expression signatures, even when the specified genes are statistically significant. Thus, we wished to verify that our identified gene lists were more biologically relevant than what we could obtain using randomly selected genes. To address these goals, we developed a process that used leave-one-dataset-out cross validation. Cross validation is a statistical method to evaluate the performance of a model on unseen data in an unbiased manner^56,57^. Rather than holding out individual samples, we held out one dataset at a time. First, we carried out a meta-analysis using *n - 1* datasets at a time (Figure 1). We assumed that if genes identified through an n - 1 meta-analysis are biologically relevant, expression values for those genes would be effective at classifying the two groups being compared on the held-out dataset. The second step was to perform classification analyses and quantify the predictive performance relative to randomly selected genes. For this step, we performed within-dataset cross validation for the held-out dataset. We used modules from the scikit-learn library (version 1.3.1)^58^ for this step. We split the datasets using the StratifiedKFold method because it ensures that each fold of the dataset has a similar proportion of observations with a given label. It is particularly useful when dealing with classification tasks that have imbalanced class distributions, as observed in our datasets. We also used the Random Forest^59^ classifier as it has been shown to perform well on gene expression data^60^. For *m* top-ranked genes (5, 10, 20, 50, 100, 500, or 1000), we performed classification and calculated a score for each cross-validation fold. We used the “balanced accuracy” scoring metric to account for the imbalanced nature of the datasets. We then repeated this process 100 times for *m* genes selected at random from the held-out dataset. We ensured that each random gene had not been identified as relevant in the meta-analysis. Consider the race datasets as an example. For each round of meta-analysis, we selected six of the seven data sets, leaving one out, and we performed the meta-analysis on those datasets. The meta-analysis produced a list of genes. From this list, we selected the top 5, 10, 20, 50, 100, 500 and 1000 genes and evaluated how accurately the expression data for those genes predicted race for the seventh data set.

**Figure 1.**
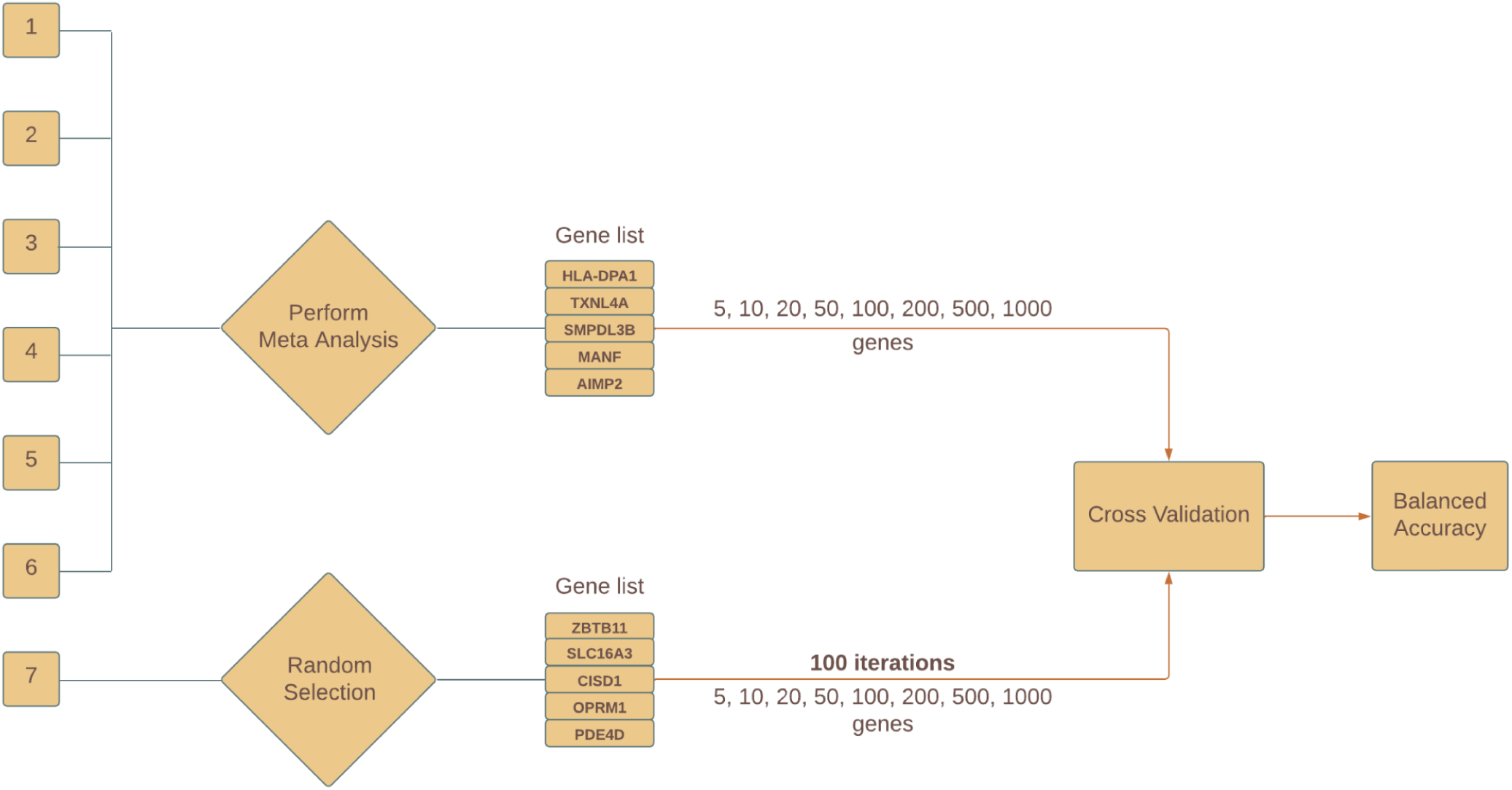
Diagram illustrating the leave-one-dataset-out cross-validation method. For a given fold, one dataset was held out for testing. Via meta-analysis, significant genes were identified and ranked according to fold change. Within-dataset cross validation was performed for the held-out test dataset, and the labels were predicted using a classification algorithm for *m* top-ranked genes. The same process was performed for randomly selected genes and repeated 100 times. We compared the balanced accuracy for the genes identified via meta-analysis vs. the genes identified via random selection.

### Pathway Analysis

Based on the cross-validation results, we investigated the gene lists using the Molecular Signatures Database (MSigDB)^61,62^ which has thousands of annotated gene sets. Examining the overlaps of these gene sets can help identify dysregulated pathways and underlying biological themes. To compute these overlaps, we pasted the top 20 genes from our gene lists into the *Investigate* tab on the MSigDB website (https://www.gsea-msigdb.org/gsea/msigdb/human/annotate.jsp). In particular, we computed overlaps using the “H: hallmark” gene sets”^63^ and the “C2: curated gene sets,” using an FDR q-value threshold of 0.05.

## Results

We performed meta-analyses to compare gene-expression levels between people of African or European descent and between people with positive or negative tumor status (ER, PR, HER2, or TNBC). There was some overlap across the analyses; some datasets had more than one of these variables (Table 1). Additionally, we compared TNBC tumors from people of African descent versus people of European descent.

**Table 1.** Dataset and sample counts for the meta-analyses. For some datasets, the original authors reported hormone receptor status but were unable to determine whether certain samples were positive or negative due to ambiguous immunohistochemistry results or when these values were marked as intermediate. These samples are listed as “Equivocal.” In other cases, the status was reported for some samples but not reported for others. These samples are listed as “Unknown.”

| Variable | Positive | Negative | Equivocal | Unknown | Total Number of Samples | Number of Datasets |
| --- | --- | --- | --- | --- | --- | --- |
|  | **White** | **Black** |  |  |  |  |
| Race | 1,175 | 295 | NA | NA | 1,470 | 7 |
| Estrogen Receptor Status | 16,174 | 4,426 | 26 | 1,334 | 21,960 | 47 |
| Progesterone Receptor Status | 11,472 | 5,359 | 96 | 1,214 | 18,141 | 35 |
| HER2/Neu Status | 2,471 | 13,429 | 179 | 1,482 | 17,561 | 28 |
|  | **Triple Negative** | **Non Triple Negative** |  |  |  |  |
| Triple Negative Status | 1,982 | 13,614 | NA | NA | 15,596 | 22 |

For the comparison between individuals of African or European ancestry, 850 genes were significantly differentially expressed (Additional File 2). For the comparisons between ER-positive and ER-negative breast tumors, 9,729 genes were differentially expressed (Additional File 3). We observed similarly high numbers for PR and HER2: 9,047 and 7,236, respectively (Additional Files 4-5). For the TNBC versus non-TNBC comparison, 8,939 genes were differentially expressed (Additional File 6). For the comparison between people of African or European ancestry specific to TNBC tumors, six genes were differentially expressed (Additional File 7).

To identify a smaller number of genes that should be considered relevant for each meta-analysis, we developed a new computational method. Leaving out one dataset at a time, we used the remaining datasets to identify significant genes and then performed classification via cross validation on the held-out dataset. We compared the predictive performance of *m* genes (as ranked by each meta-analysis) against the predictive performance of randomly selected genes, to estimate the number of significant genes that would be most predictive. This analysis revealed that 20 genes often performed well compared to other thresholds (Figures 2-6). Although this threshold did not always perform better than other thresholds, we chose 20 genes as the threshold for all six meta-analyses for consistency. Twenty genes is few enough that researchers can interpret such a list and potentially create a panel to use in a laboratory setting. Table 2 shows the top 20 genes for each comparison. *GATA3, CA12, XBP1* and *FOXA1* were among the top 20 genes for ER, PR, and TNBC. However, none of these overlapped with the genes for HER2 status (or race).

**Figure 2.**
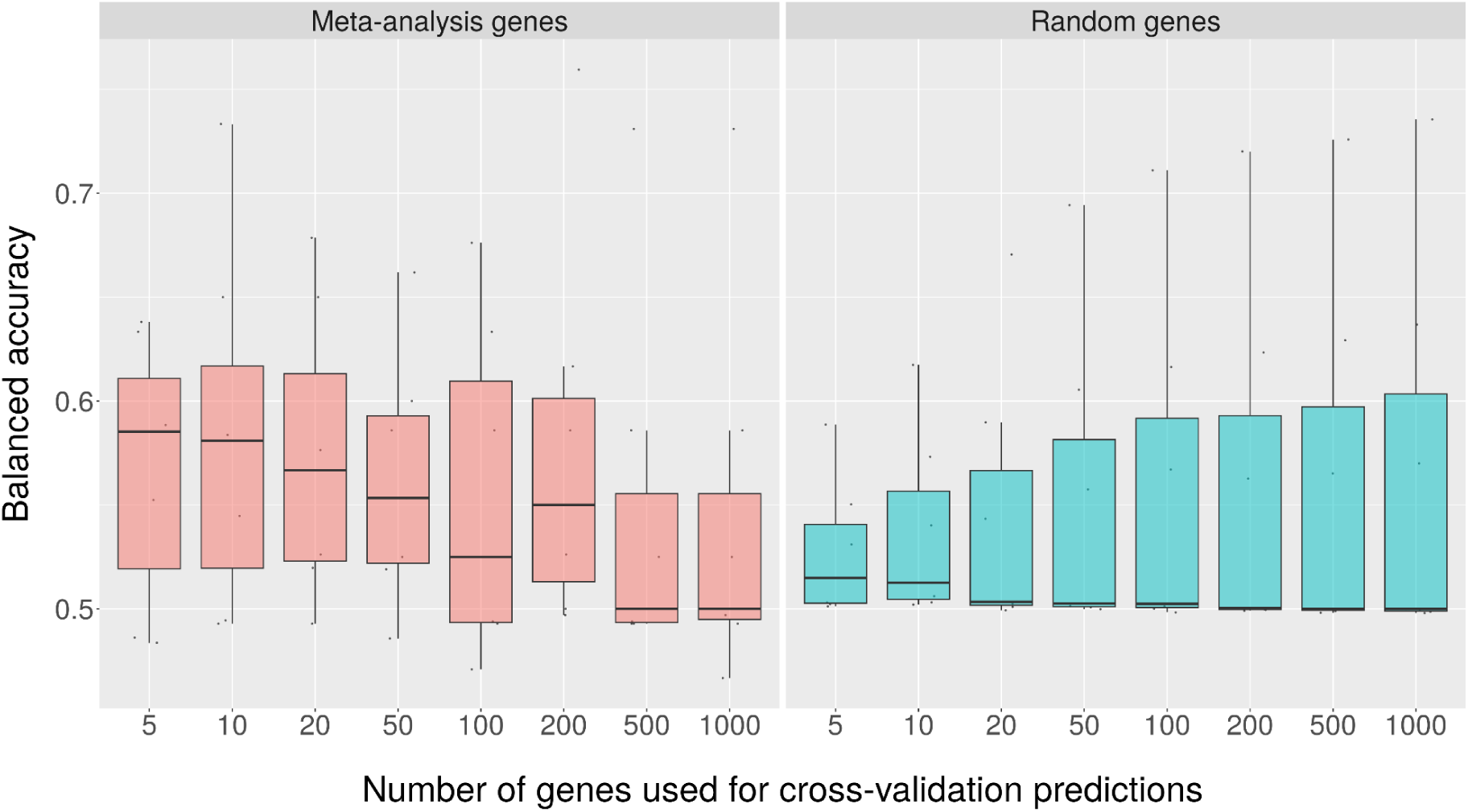
Cross-validation results for race status. The left panel represents the balanced accuracy, identified via cross validation on each held-out test set, for different numbers of top-ranked genes from the race meta-analysis. Each data point represents the balanced accuracy for a particular dataset. The right panel represents the balanced accuracy for the same numbers of genes selected at random in 100 iterations. Each data point represents the mean balanced accuracy for a particular dataset, across the iterations. The balanced accuracy for the meta-analysis genes was relatively low (between 0.5 and 0.6) when using 5 to 200 genes; it was equivalent to baseline levels (0.5) when 500 or 1000 genes were used. The balanced accuracy for the randomly selected genes was consistently close to baseline levels.

**Table 2.** Top-ranked genes identified in the meta-analyses. . These genes represent our findings when performing meta-analyses across all datasets for which each variable was available (i.e., not as part of our cross-validation experiments). Fewer than 20 genes are shown for “Race + TNBC” because this meta-analysis identified only 6 genes with statistical significance. Genes bolded and highlighted in red were common across more than one of the meta-analyses.

| Rank | Race | ER Status | PR Status | Her2 Status | TNBC | Race + TNBC |
| --- | --- | --- | --- | --- | --- | --- |
| 1 | DNAJC15 | **GATA3** | PGR | ERBB2 | **FOXA1** | RETN |
| 2 | HLA-DPA1 | ESR1 | **CA12** | PGAP3 | MLPH | ADAMTS13 |
| 3 | STAP2 | **CA12** | MAPT | STARD3 | FOXC1 | ACOXL |
| 4 | CEP68 | TBC1D9 | **GATA3** | GRB7 | **GATA3** | FAM3A |
| 5 | MOGS | **FOXA1** | ESR1 | CDK12 | EN1 | GPR35 |
| 6 | TNS1 | MLPH | DNALI1 | PSMD3 | GABRP | PGLYRP1 |
| 7 | TXNL4A | DNALI1 | GREB1 | MED1 | **CA12** |  |
| 8 | PSPH | NAT1 | NAT1 | GSDMB | BCL11A |  |
| 9 | EIF5A | PSAT1 | SCUBE2 | MED24 | SFT2D2 |  |
| 10 | RPP25 | SLC39A6 | IL6ST | PNMT | **XBP1** |  |
| 11 | PKP3 | **XBP1** | TBC1D9 | TCAP | AGR2 |  |
| 12 | POP7 | ABAT | ABAT | MSL1 | FAM171A1 |  |
| 13 | CENPA | GREB1 | GFRA1 | CASC3 | DSC2 |  |
| 14 | TTBK2 | MAPT | **XBP1** | CWC25 | TBC1D9 |  |
| 15 | CXCL12 | THSD4 | KDM4B | PSMB3 | UGT8 |  |
| 16 | PYCR1 | BCL11A | THSD4 | PIP4K2B | SPDEF |  |
| 17 | HSD17B10 | GFRA1 | CELSR1 | LASP1 | PSAT1 |  |
| 18 | TIMM23 | AGR2 | SLC39A6 | RPL23 | SOX10 |  |
| 19 | PCGF1 | PHGDH | **FOXA1** | CLCA2 | ART3 |  |
| 20 | ABCA6 | DSC2 | ERBB4 | KMO | MICALL1 |  |

For the comparison between tumors from individuals of European or African descent, the median balanced accuracy for the top 20 genes was 0.57 (the maximum possible accuracy is 1.0). The median balanced accuracy for the randomly selected genes was 0.50. Using a one-sided, Mann-Whitney U test, we compared the balanced accuracy values when using the top 20 meta-analysis genes for classification, versus the balanced accuracies for the randomly selected genes; the meta-analysis accuracies were *not* significantly higher (p-value = 0.23; Figure 2). These findings suggest that gene-expression differences between these populations were not significantly different from randomly selected genes; however, sample sizes may have been a limiting factor.

For the ER-positive versus ER-negative comparison, the balanced accuracies were significantly higher for the meta-analysis genes than for the randomly selected genes (p < 0.001; Figure 3). The same was true for the PR, HER2, and TNBC comparisons (p < 0.001).

**Figure 3.**
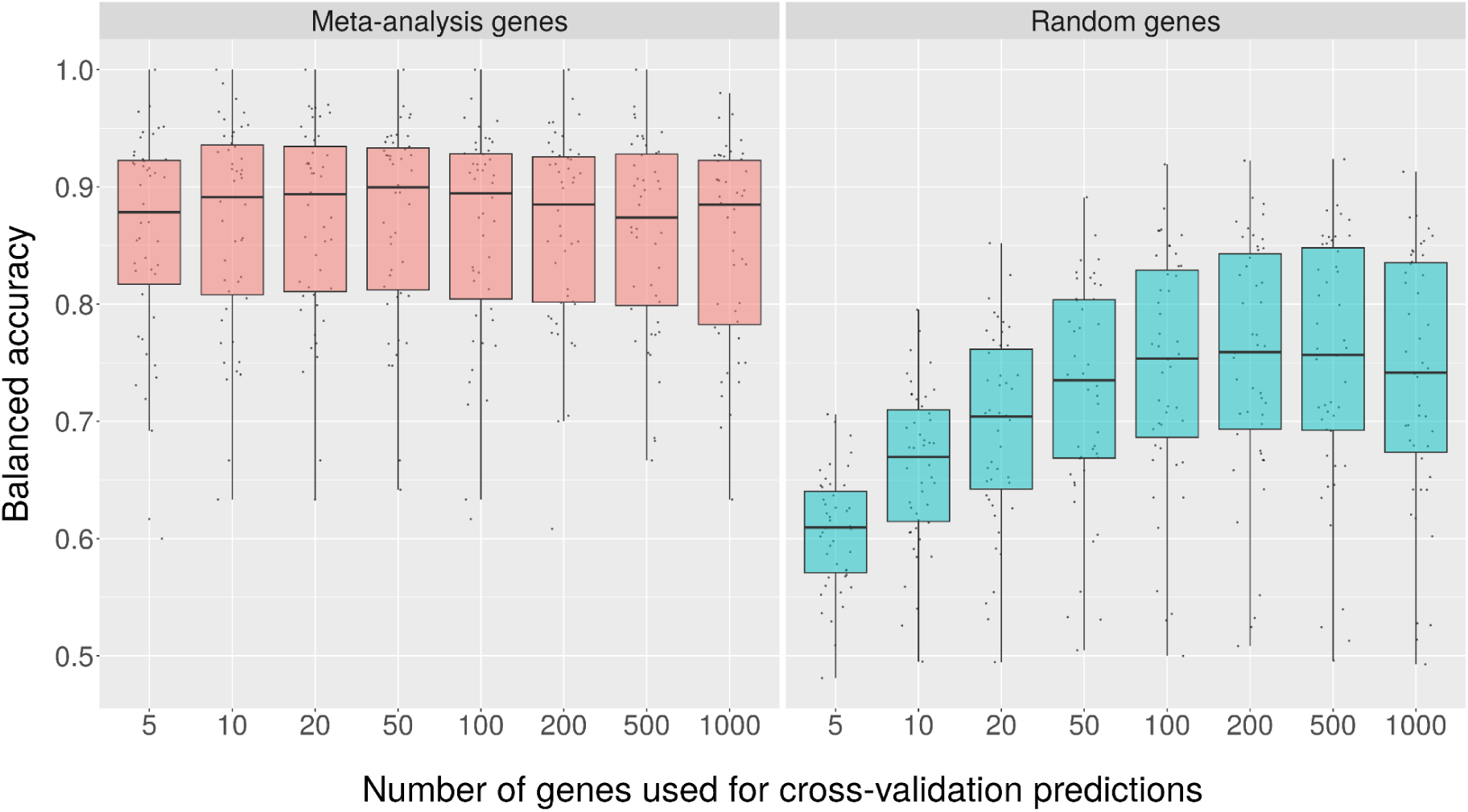
Cross-validation results for ER Status. The left panel represents the balanced accuracy, identified via cross validation on each held-out test set, for different numbers of top-rank genes from the ER meta-analysis. Each data point represents the balanced accuracy for a particular dataset. The right panel represents the average balanced accuracy for the same numbers of genes selected at random in 100 iterations. Each data point represents the mean balanced accuracy for a particular dataset, across the iterations. The balanced accuracy for the meta-analysis genes was consistently high (around 0.85 to 0.9) regardless of the number of genes used, while the balanced accuracy for the random genes started low and increased as the number of genes increased.

However, the overall patterns were markedly distinct for HER2 relative to the other comparisons (Figures 3-6). For nearly all thresholds, the balanced accuracy was above 0.8 when using the meta-analysis genes; however, for ER, PR, and TNBC, the random-gene accuracies started low (around 0.6) and increased considerably as the number of genes increased. This pattern suggests that ER and/or PR positivity cause a cascade of downstream effects, possibly altering the expression of thousands of genes. Thus, when we selected genes at random (not including the top-ranked genes), the classification algorithm was frequently able to classify the tumors as positive or negative. In contrast, for HER2, the median balanced accuracy was below 0.55 for all thresholds when randomly selected genes were used. This pattern suggests that overexpression of HER2 protein, which typically results from DNA amplification or somatic mutations^64^, thus activating oncogenic pathways^65,66^, causes relatively few downstream effects compared to endocrine breast cancers, which rely on estrogen or progesterone signaling for tumor growth^67,68^.

**Figure 4.**
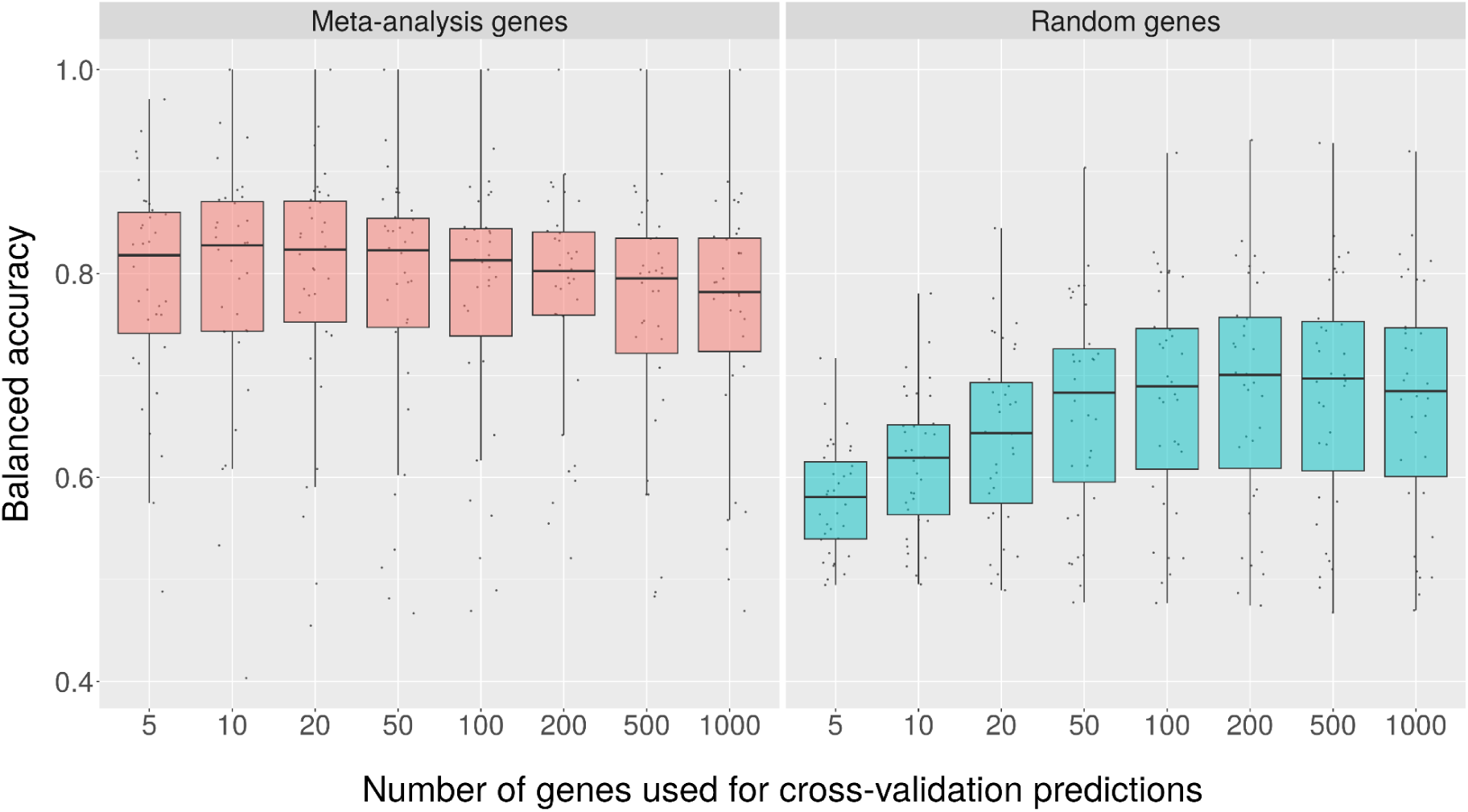
Cross-validation results for PR Status. The left panel represents the balanced accuracy, identified via cross validation on each held-out test set, for different numbers of top-rank genes from the PR meta-analysis. Each data point represents the balanced accuracy for a particular dataset. The right panel represents the average balanced accuracy for the same numbers of genes selected at random in 100 iterations. Each data point represents the mean balanced accuracy for a particular dataset, across the iterations. The balanced accuracy for the meta-analysis genes was consistently high (around 0.8) regardless of the number of genes used, while the balanced accuracy for the random genes started low and increased as the number of genes increased.

**Figure 5.**
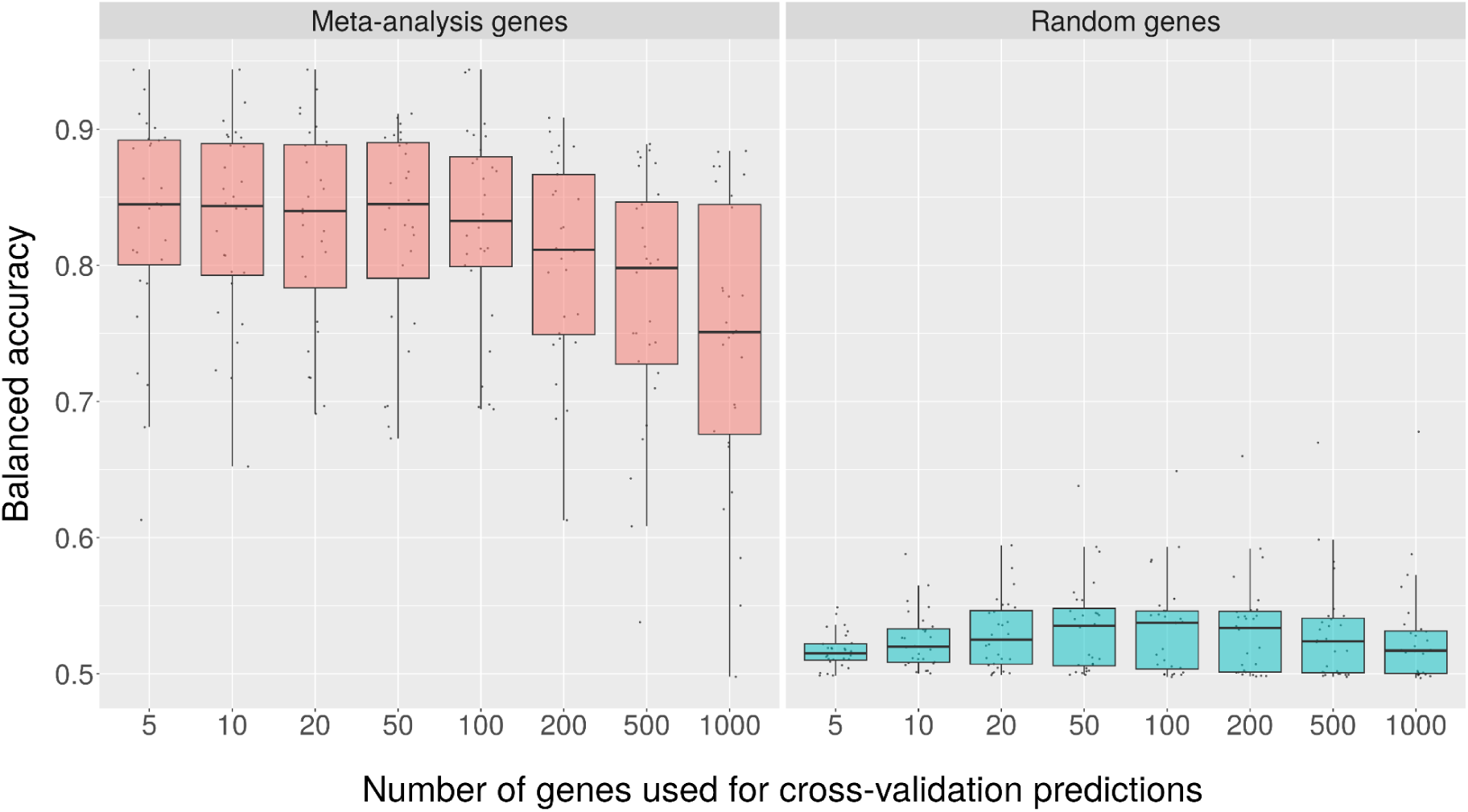
Cross-validation results for HER2 status. The left panel represents the balanced accuracy, identified via cross validation on each held-out test set, for different numbers of top-rank genes from the HER2 meta-analysis. Each data point represents the balanced accuracy for a particular dataset. The right panel represents the average balanced accuracy for the same numbers of genes selected at random in 100 iterations. Each data point represents the mean balanced accuracy for a particular dataset, across the iterations. The balanced accuracy for the meta-analysis genes was consistently higher than the balanced accuracy for the random genes.

**Figure 6.**
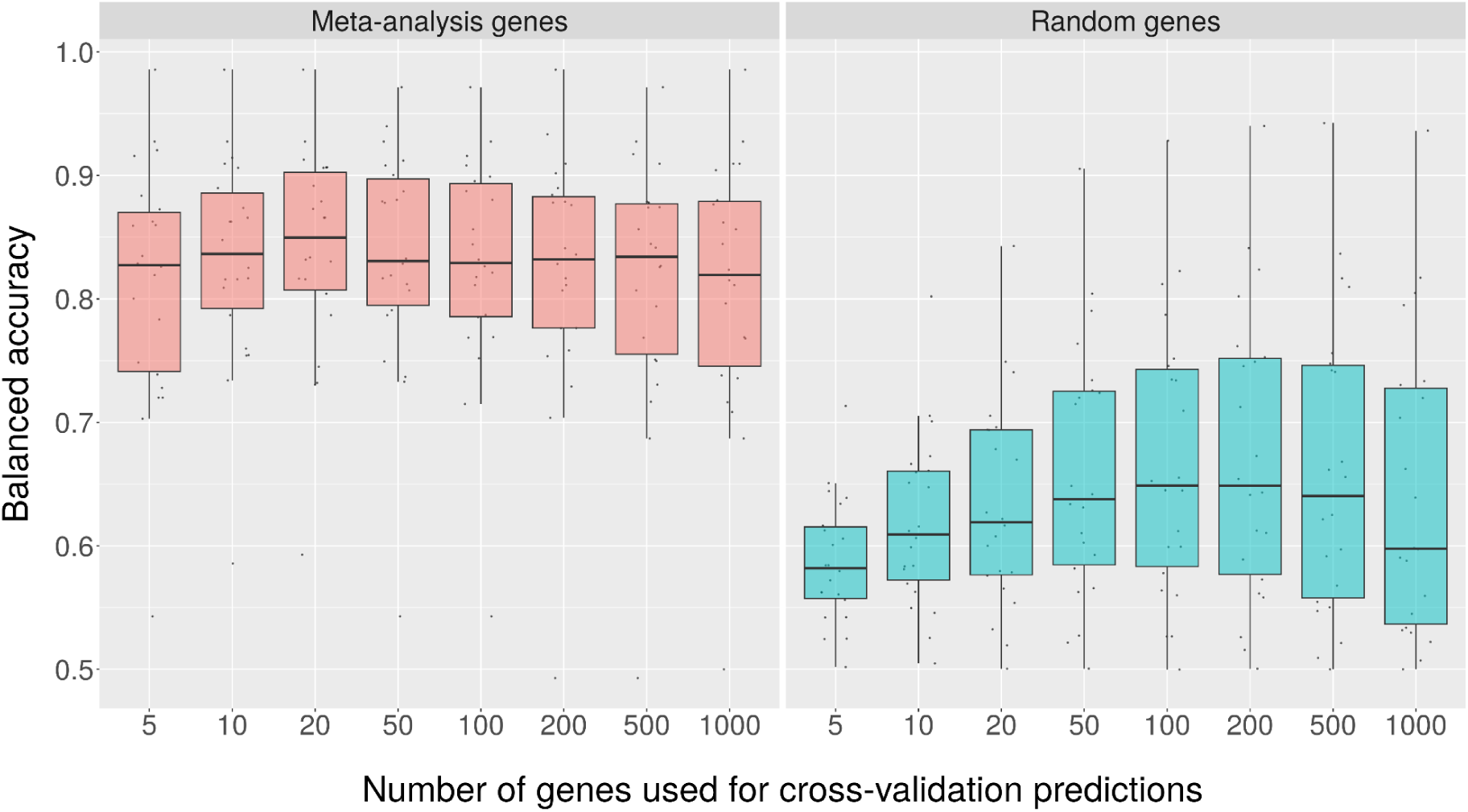
Cross-validation results for TNBC status. The left panel represents the balanced accuracy, identified via cross validation on each held-out test set, for different numbers of top-rank genes from the TNBC meta-analysis. Each data point represents the balanced accuracy for a particular dataset. The right panel represents the average balanced accuracy for the same numbers of genes selected at random in 100 iterations. Each data point represents the mean balanced accuracy for a particular dataset, across the iterations. The balanced accuracy for the meta-analysis genes was consistently higher than the balanced accuracy for the random genes.

For the comparison between individuals of African or European descent, some of the top genes were *DNAJC15*, *RPP25*, *CXCL12*, *EIF5A*, and *TIMM23*. These genes play diverse roles in cellular functioning. *DNAJC15* has been found to be expressed in breast cancer cells^69^ and may play a role in apoptosis^70^. *RPP25* enables ribonuclease P RNA binding activity and is involved in tRNA 5’-leader removal^71,72^. *CXCL12* is the ligand for the G-protein coupled receptor and chemokine and can affect tumor progression, angiogenesis, metastasis, and survival in cancer cells^73,74^. *EIF5A* is a translation elongation factor^75^ and plays a role in regulating tumor cell growth^76^. *TIMM23* mediates the transport of transit peptide-containing proteins across the membrane^77^. Elvidge et al. showed that some of these genes may play a role in modulating changes in gene expression induced by hypoxia in MCF-7 breast cancer cell lines^78^. However, hypoxia and these other cellular processes do not have a clear, known function that would explain gene-expression differences in breast tumors between people of African or European descent specifically.

The top genes identified in the ER-positive versus ER-negative comparison were *GATA3*, *ESR1,* and *CA12*. These genes exhibited approximately two-fold higher expression in ER-positive patients compared to ER-negative patients. *ESR1* encodes the estrogen receptor alpha (ERα) protein, which mediates the effects of estrogen in various tissues throughout the body^79^. Finding this gene near the top of our list when comparing ER positive versus ER negative breast cancers is evidence of the validity of our data and methods. *GATA3* is crucial for the differentiation of the mammary epithelium and is a key regulator in breast cancer^80^. It also plays a role in mediating the binding of estrogen receptor elements to *ESR1*^81^. Furthermore, co-expression of *GATA3*, *ESR1,* and *FOXA1* in breast tumors has been reported^82,83^. *CA12* is involved in cellular proliferation and pH regulation and has been associated with a good prognosis in breast cancer^84^. Other significant genes include *FOXA1* and *TBC1D9*. *FOXA1* is a transcriptional regulator that modulates the effects of estrogen signaling by influencing *ESR1* activity^85,86^. *TBC1D9* is involved in cellular trafficking processes that can influence cancer cell behavior^87^. These and other top-ranked genes merit further study to provide mechanistic insight into the ER protein’s role in breast tumorigenesis, and suggest their potential roles both as biomarkers and therapeutic targets.

For the comparison between PR-positive and PR-negative tumors, a common thread among the top genes identified, is their involvement in various aspects of known breast cancer biology. Other studies^88^ have shown that many of these genes (*PGR, ESR1, FOXA1, GATA3, GREB1*) are differentially expressed and that *PGR*^89^ and *CA12*^84^ may be good prognostic markers for breast cancer.

Among the top genes for the comparison between HER2-positive and HER2-negative tumors were *ERBB2* (which was expected because this gene encodes the HER2 protein) as well as *PGAP3, STARD3, GRB7, CDK12,* and *PSMD3. PGAP3* is overexpressed in breast cancer patients, is associated with lymph node metastasis, and correlates with HER2 overexpression^90^. *STARD3* is necessary for the transfer of cholesterol and metabolism in tumor cells^91^, and is often co-amplified with *HER2*^92^. *GRB7* is responsible for signal transduction from cell surface receptors via signaling pathways in the cell^93^. It is sometimes co-amplified and overexpressed with HER2 and is associated with a more aggressive phenotype^94^. *CDK12* is a transcription-associated cyclin-dependent kinase, which plays various roles in the regulation of translation^95^, cellular proliferation^96^, and cell cycle progression^97^; it acts as a tumor promoter in HER2-positive breast cancer^98^. *PSMD3* is an enzyme regulator by stabilizing *HER2* and preventing it from degradation^99^. These genes may serve as alternative markers for HER2 positive breast cancer^90,100^.

For the comparison between TNBC and non-TNBC patients, the most significant genes sometimes overlapped with what we identified in the earlier analyses. *FOXA1*, *MLPH*, and *GATA3* were expressed at approximately two-fold higher levels in non-TNBCs compared to TNBCs, while *FOXC1* had approximately 2-fold lower expression in non-TNBC than TNBC tumors.

Our last comparison focused on individuals of African or European ancestry who had TNBC. The statistically significant genes were *RETN, ADAMTS13, ACOXL, FAM3A, GPR35*, and *PGLYRP1*. *RETN* encodes the hormone resistin that plays a role in inflammation^101^ and insulin resistance^102^, which are critical factors in obesity-related cancers^103^. Increased resistin levels have been associated with breast cancer^104,105^. *ADAMTS13* encodes an enzyme that processes von Willebrand factor, a protein involved in blood clotting. Dysregulation of *ADAMTS13* has been implicated in cancer metastasis and tumor angiogenesis due to its role in extracellular matrix remodeling^106,107^. *ACOXL* is involved in fatty acid β-oxidation and metabolic processes^108^. Although there is no research linking it to breast cancer, its expression has been linked to prostate cancer, where it is a potential biomarker due to differential expression in benign versus malignant prostate tissues^109^. *FAM3A* is involved in regulating of both glucose and lipid metabolism^110^. Given its metabolic functions and the link between metabolism and cancer progression, it has been suggested to play a role in metabolic diseases^111^ and possibly cancers^112^. *GPR35* is involved in inflammatory responses and is highly expressed in various cancers, including gastrointestinal and prostate. It is considered a potential therapeutic target due to its role in tumorigenesis and cancer progression^113,114^. *PGLYRP1* encodes a protein involved in the innate immune response and plays a role in regulating acquired immunity^115^. *PGLYRP1* has been implicated in inflammation and immune responses in the tumor microenvironment^116,117^, which can influence cancer development and progression. These genes share involvement in inflammation, metabolic regulation, and the tumor microenvironment, which are critical processes in the development and progression of cancer even though they are not all specific to breast cancer. We did not find evidence to provide mechanistic insight about these expression differences between people of African or European descent. Yet they may merit further exploration as potential therapeutic targets.

To gain further insight into the biology behind the selected genes for each comparison, we performed pathway enrichment analysis using MSigDB^61,62^ (Additional Files 8 - 12). Some of these results further confirm the validity of our methods. For example, in the ER-positive versus ER-negative comparison, curated pathways that reflect ER signaling were statistically significant. Similarly, for the HER2-positive versus HER2-negative comparison, pathways associated with HER2 amplification were significant. For the PR comparison, ER pathways were statistically significant, but we did not observe pathways specific to PR signaling; this suggests that ER and PR positivity have similar effects but may also reflect that MSigDB lacks pathways specific to PR. When a gene is statistically significant but not associated with any of the statistically significant pathways for a given comparison, it serves as a hypothesis for further investigation. Additionally, for significant pathways, genes in those pathways might be prioritized for investigation, even when they were not among the most statistically significant genes.

## Discussion

Researchers have used diverse methodologies to analyze and combine gene expression datasets^118^. Some have focused on identifying prognostic markers^43,119^ or genes^120,121^, predicting treatment outcomes^122–124^, or deriving more specific breast cancer subtypes^49,125^. Others have attempted to differentiate between primary and metastatic tumors^126^, as well as differential gene-expression analysis^43,127^. In this study, we carried out meta-analyses for tens of thousands of breast tumors. To the best of our knowledge, no meta-analyses of this magnitude has been performed previously; nor has a meta-analysis been performed to compare breast cancer patients of African and European ancestry. Schneider et al. focused on ER patients and identified a subset of genes that also show up in our analysis^128^. Al-Ejeh et al. focused on TNBCs and identified a set of 206 “aggressive genes”^30^, some of which also were significant in our study. D’Arcy et al. compared Luminal A breast tumors between African American and Caucasian women and arrived at a list of 6 genes; but only one of those genes (*PSPH*) overlaps with our list. These differences may be due to their modeling approach and the fact that they had a smaller sample size (n = 167 samples) and used an imputation method to account for missing data.

### Race and Gene Expression

One goal of this research was to understand the biology behind disparities in breast cancer between women of African and Caucasian descent. Our analysis revealed 850 genes that were significantly differentially expressed between these populations. Among the top 20 genes, we observed significant roles for inflammation (*RETN*)^101^, cell cycle regulation (*CENPA* and *TTBK2*)^129,130^, immune responses (*HLA-DPA1, CXCL12,* and *STAP2*)^74,131,132^, and cellular metabolism (*PSPH, PYCR1,* and *HSD17B10*)^133–135^. These can influence tumor behavior and responses to treatments. For instance, *RETN* (resistin) is associated with inflammation and insulin resistance, which are critical in the pathogenesis of obesity-related cancers, including breast cancer. Elevated resistin levels have been linked to poor prognosis in breast cancer patients, underscoring the potential of RETN as a biomarker for breast cancer in diverse populations.

Black women are more likely to be diagnosed with TNBC, which is more aggressive and has fewer targeted treatment options, while Caucasian women are more frequently diagnosed with hormone receptor-positive breast cancers, which have better prognoses and more effective hormone-targeted therapies^136,137^. The genes we identified when we merged TNBC and race are involved in inflammation and immune responses, which are critical pathways in cancer development, including breast cancer. ADAMTS13, an enzyme involved in blood clotting, plays a role in cancer metastasis and angiogenesis through extracellular matrix remodeling. Dysregulation of ADAMTS13 has been implicated in various cancers, suggesting its potential as a therapeutic target in breast cancer^138^. Similarly, genes such as *GPR35*, involved in inflammatory responses, and *PGLYRP1*, implicated in the innate immune response, highlight the importance of inflammation and immune regulation in cancer development^139–143^. Variations in these genes and their expression can vary by race, potentially influencing disease susceptibility and progression^144,145^.

A key limitation of our analysis is that categories based on race and ethnicity are self-described and may not adequately represent a person’s ancestry. By performing secondary analyses of publicly available data, we were limited to the descriptions provided by the original authors. For simplicity, we used broad groupings (African descent, European descent) to categorize the patients. However, by aggregating data from many studies, we anticipated that meaningful patterns would be identifiable, despite this noise. Although our analyses identified genes that have plausible connections to breast tumorigenesis and may lead to insights about population disparities, gene-expression differences between these categories were relatively minor, providing support for the argument that race is a social construct more than a biological concept^146^.

### ER, PR, and HER2 Status

Compared to ER-negative breast tumors, genes like *ESR1, GATA3,* and *CA12* were more highly expressed in ER-positive tumors. *ESR1*, encoding the estrogen receptor alpha (ERα), is crucial for mediating estrogen’s effects and is a well-established marker in breast cancer diagnostics and treatment^79,147^. *GATA3* is essential for mammary epithelial differentiation and is a key regulator in breast cancer, while *CA12* is involved in cellular proliferation and pH regulation and is associated with a good prognosis^148,149^.

For PR status, genes such as *PGR, ESR1,* and *FOXA1* were differentially expressed, consistent with their roles in breast cancer biology. Progesterone receptor (PGR) and estrogen receptor (ESR1) are critical markers for hormone receptor-positive breast cancers and are important for determining therapeutic strategies^150,151^.

Our HER2-positive vs. HER2-negative comparisons identified genes like *ERBB2*, *PGAP3*, and *GRB7*, which were prominently expressed in HER2-positive tumors. *ERBB2* overexpression is a hallmark of HER2-positive breast cancers and a target for therapies such as trastuzumab^152^. The co-amplification of genes like *GRB7* with HER2 suggests their roles in aggressive breast cancer phenotypes and potential as additional therapeutic targets^153,154^.

### TNBC vs. Non-TNBC

TNBC vs. non-TNBC analysis identified *FOXA1, MLPH,* and *GATA3,* which showed higher expression in non-TNBC, while *FOXC1* was expressed lower in non-TNBC. These findings highlight the distinct molecular profiles of TNBC, characterized by the lack of hormone receptor and HER2 expression^155^.

### Common Genes Across Comparisons

*GATA3, CA12, TBC1D9, XBP1,* and *FOXA1* were consistently significant across ER, PR, and TNBC comparisons but not in the HER2 comparison. This supports the notion that the mechanisms behind ER-positive and PR-positive tumors are overlapping and that HER2-positive tumors are mechanistically distinct from endocrine breast cancers, necessitating different therapeutic strategies^156–158^.

### Computational Validation

It is common for differential-expression studies, such as this meta-analysis, to identify thousands of statistically significant genes. Such findings may be reassuring because they imply that the compared phenotypes are truly distinct at the molecular level. However, a larger goal of a differential-expression study is to reduce the search space for genes that are most important in driving the phenotype. Lab-based, experimental validation can provide assurances that genes are truly differentially expressed between conditions; however, such procedures do not scale beyond a few genes. Furthermore, the complexity of cancers likely cannot be captured adequately using data from a few genes. Therefore, it is desirable to find a balance between characterizing the complexity of cancers and identifying a gene set that is small enough to be analyzed and interpreted by humans and that can be explored feasibly in subsequent, mechanistic studies. Our novel computational method addresses these goals, while also showing that most of our identified gene sets are more relevant than randomly selected genes. Moreover, it provides insight into the potential to use these genes to predict phenotypes. A limitation of differential-expression studies is that commonly used statistical methods consider evidence from one gene at a time, whereas our classification approach constructs models from many genes simultaneously. We believe this method has potential to be applied in many other contexts. Further validation will be necessary.

## Conclusion

Findings from our meta-analyses elucidate gene expression differences between racial groups and clinically relevant breast cancer subtypes, highlighting the roles of diverse cellular processes. They provide additional evidence about mechanisms that may drive breast cancer biology. Such studies are necessary to pave the way for developing targeted diagnostics and therapies in a personalized-medicine context. Further research is warranted to explore the mechanistic pathways of these genes and their potential as biomarkers and therapeutic targets in diverse patient populations.

## List of Abbreviations

ER: estrogen receptor
ESR1: estrogen receptor
HER2: human epidermal growth factor receptor 2
IGF1: insulin-like growth factor 1
IHC: immunohistochemistry
MSigDB: Molecular Signatures Database
OSF: Open Science Framework
PGR: Progesterone receptor
PR: progesterone receptor
SCAN-B: The Sweden Cancerome Analysis Network - Breast
TCGA: The Cancer Genome Atlas
TNBC: Triple-negative breast cancers

## Declarations

### Ethics approval and consent to participate

Not applicable

### Consent for publication

Not applicable

### Availability of data and materials

All source code, gene expression data, and metadata are available in a publicly accessible Open Science Framework (OSF)^159^ repository (https://osf.io/mr8sb/). The code is packaged and executable using Docker^160^ software containers for easier reproducibility.

### Authors’ contributions

ION prepared and analyzed data. Both authors designed the study and interpreted the results. SRP supervised the work. Both authors wrote the manuscript and approved the final manuscript.

## Acknowledgements

Not applicable

